# Extrusion-Printed Silicone Microarchitectures for Geometry-Controlled Flow in Lateral Flow Diagnostics and Paper Microfluidics

**DOI:** 10.64898/2026.05.19.726334

**Authors:** Mecit Altan Alioglu, Satheesh Natarajan, David Skrodzki, Oguzhan Colak, Dipanjan Pan

**Author notes:** Corresponding author at: The Pennsylvania State University, University Park, PA, 16802, USA. E-mail addresses.

## Abstract

Paper-based diagnostics such as lateral flow assays (LFAs) and microfluidic paper-based analytical devices (µPADs) have attracted considerable attention because of their low cost, portability, and ease of use. Currently, to enable fabrication of µPADs and improve LFA performance, hydrophobic blocks are patterned on paper substrates. However, fabrication of high-resolution hydrophobic barriers remains a major challenge. In this work, we developed a novel silicone extrudable ink for the fabrication of hydrophobic features on paper substrates. The ink was formulated using a vinyl-terminated polydimethylsiloxane (vPDMS) and polymethylhydrosiloxane (PMHS) system crosslinked through platinum-catalyzed hydrosilylation, and its rheological properties were tailored by incorporating silica fillers, obtaining a shear-thinning gel suitable for extrusion. The resulting formulation provided tunable properties, controlled deposition, and stable feature formation, enabling simple, low-cost, rapid, and robust fabrication of high-resolution hydrophobic barriers. Using this approach, we demonstrated improved fluid confinement and pattern fidelity on paper substrates, fabricated high-resolution paper microfluidic devices down to 150 µm channel width, and enhanced the sensitivity of an LFA for a malaria diagnostic test. These results highlight the potential of this silicone ink platform as a practical and scalable strategy for advancing high-performance paper-based diagnostic technologies.

## 1. Introduction

Accurate and accessible diagnostic tools are essential for modern healthcare because they enable early disease detection, guide treatment decisions, monitor disease progression, and improve patient outcomes while reducing overall healthcare costs. In addition to their role in clinical settings, diagnostics are also critical for public health surveillance, outbreak control, and health management in resource-limited environments [1– 3]. Among the many diagnostic platforms available, lateral flow assays (LFAs) have attracted significant attention due to their simplicity, low cost, rapid response, and ease of use without the need for complex instrumentation or highly trained personnel [4,5]. These paper-based devices are widely used for point-of-care testing because they can provide fast and user-friendly detection of a broad range of analytes, including pathogens, biomarkers, and small molecules. Due to these advantages, LFAs have become an important tool for decentralized diagnostics and continue to be developed for improved sensitivity, reliability, and broader applications [6,7].

In addition to the widespread use of LFAs, the broader field of paper microfluidics has emerged as an important area of research for developing simple, low-cost, and accessible analytical devices. Although paper microfluidic systems share fundamental principles of capillary-driven flow and porous substrate-based transport with LFAs, they extend beyond conventional test-strip formats and offer greater flexibility in device architecture and fluid manipulation [8,9]. Conventional microfluidics has enabled precise control and manipulation of very small fluid volumes, supporting advanced functions such as mixing, separation, and multiplexed analysis in compact devices [10]. Building on these principles, microfluidic paper-based analytical devices (µPADs) have established themselves as a versatile platform by combining fluid handling capabilities with the affordability, simplicity, and accessibility of paper-based materials. In paper microfluidic systems, fluids move through porous substrates by capillary action, eliminating the need for external pumps and making the devices easy to operate [11,12]. The low cost of paper, ease of fabrication, compatibility with biochemical assays, and ability to handle small sample volumes make paper microfluidics especially attractive for point-of-care testing and use in resource-limited settings. Paper microfluidics is steadily improving and is expected to play an even greater role in future diagnostic technologies [13–15].

One of the major strategies for improving both the performance of LFAs and the fabrication of µPADs has been the integration of hydrophobic barriers into paper substrates [7,13,16]. These hydrophobic features enable controlled fluid transport by defining flow paths, limiting lateral spreading, and separating functional regions within the device. As a result, they can improve assay performance, enhance signal localization, and enable the development of more complex paper-based microfluidic designs. Because of these advantages, the formation of hydrophobic barriers has become a key aspect of paper microfluidic device engineering and optimization [14,17]. Currently, a variety of techniques have been used to introduce hydrophobic features into paper substrates, including photolithography, wax patterning, wax dipping, stamping, and inkjet printing [17–22]. These methods have played an important role in the development of paper microfluidic devices by enabling controlled fluid pathways and functional device architectures. However, despite their widespread use, many of these approaches face limitations when high-resolution channel fabrication is required. In particular, the diffusion or spreading of hydrophobic materials within the porous paper matrix can reduce pattern fidelity and make it difficult to fabricate channels with high resolution. Although with some of these methods can achieve higher resolution they have to rely on specialized instrumentation, multistep processing, or costly fabrication setups [14,17].

Malaria remains a major global health challenge, especially in resource-limited regions where rapid and accurate diagnosis is essential [23]. Plasmodium lactate dehydrogenase (pLDH) is a reliable biomarker because it is expressed by all human malaria parasites during active infection and reflects the presence of viable parasites [24]. Unlike HRP2, which is specific to P. falciparum and can persist after parasite clearance, pLDH correlates more closely with active infection and treatment response [25]. This is particularly important in regions with HRP2/HRP3 gene deletions, where HRP2-based assays show reduced diagnostic performance [26]. Therefore, pLDH is increasingly recognized as a preferred biomarker for next-generation point-of-care malaria diagnostics. Due to these advantages, we selected pLDH as the target biomarker to demonstrate the applicability of our lateral flow assay platform.

Here we utilized a novel technique to manufacture hydrophobic blocks using a silicone extrudable ink. This approach offers several important advantages, including simple processing, low cost, robustness, rapid fabrication, and the ability to generate high-resolution features. By enabling controlled deposition of the silicone ink onto the paper substrate, the method minimizes undesired spreading and allows the formation of sharper and more well-defined hydrophobic barriers. As a result, it improves pattern fidelity and resolution while avoiding the need for complex fabrication steps or expensive instrumentation. These advantages make the technique highly attractive for the scalable manufacturing of paper-based microfluidic devices with improved fluid control and device performance. Silicone was selected as the barrier material because of its chemical and thermal inertness, mechanical durability, and long-term stability under a wide range of environmental conditions [27]. Although polydimethylsiloxane (PDMS) has previously been used to pattern hydrophobic blocks, the viscous liquid nature of unmodified commercial PDMS resulted in spreading and limited patterning resolution [28–30].

To obtain an extrudable silicone ink, we formulated a vinyl-terminated polydimethylsiloxane (vPDMS) and polymethylhydrosiloxane (PMHS) mixture that undergoes platinum-catalyzed hydrosilylation crosslinking. To achieve the rheological properties required for extrusion-based 3D printing, the ink was modified by incorporating hydrophobically treated silica fillers together with a thixotropic agent, resulting in a shear-thinning gel suitable for controlled deposition. This custom formulation provides tunable material properties and enables extrusion with shape retention after printing. To demonstrate the capabilities of this approach, we used the formulated ink to improve the sensitivity of a lateral flow assay for malaria diagnostic test and to fabricate paper microfluidic devices with high-resolution and complex channel features. Overall, this technique is a promising strategy for the fabrication of advanced LFAs and paper microfluidic devices.

## 2. Materials and Methods

### 2.1. Materials

Vinyl terminated polydimethylsiloxane (vPDMS) (433012), Polymethylhydrosiloxane (PMHS) (176206), Platinum(0)-1,3-divinyl-1,1,3,3-tetramethyldisiloxane complex (Karstedt’s catalyst) (479519), 1-Ethynyl-1-cyclohexanol (E51406), NC membrane (HF-180 PLUS) (Merck Millipore). 10x PBS buffer, 10% BSA from Thermo-Fischer, 20nm Gold NP and Tween-20 were acquired from Sigma–Aldrich (St. Louis, MO, USA). THI-VEX was acquired from Smooth-On (Macungie, PA, USA). Dimethyldicholorosilane treated fumed silica filler (SF) (HD-162) was acquired from Tunoff (China). Sample pad and absorbent pad were acquired from Cytiva (Marlborough, MA, United States). Detection Mouse anti-pLDH Pan Malaria IgG antibody, Antigen-*Pan Malaria* pLDH, Capture Mouse anti-pLDH Pan Malaria IgG antibody, anti-mouse IgG were acquired from Vista laboratory services (Langley, WA, USA).

### 2.2. Preparation of extrudable hydrophobic silicone ink

The extrudable hydrophobic ink was prepared using a SpeedMixer DAC-330-110 SE (Flacktek Manufacturing, Landrum, SC, USA). The preparation of a 5 g 8:2 vPDMS:PMHS formulation with 8% Si content was carried out as follows. First, 3.64 g vPDMS, 5 µl Karstedt’s catalyst and 1 µl 1-Ethynyl-1-cyclohexanol is placed in a 12 ml container and mixed at 3500 rpm for 30 sec. Then, 0.4 g SF is included in the container and mixed at 3500 rpm for 5 min. Then, 0.05 g thixotropic agent (THI-VEX) is included in the container and mixed at 3500 rpm for 3 min. Following the mixing, the mixture was then allowed to rest for 15 min to equilibrate to room temperature. Finally, 0.93 g of PHMS was included to the container and mixed at 3500 rpm for 3 min. After mixing was completed, the ink is loaded into a 3 or 5 ml syringe barrel and centrifuged at 4000 rpm for 4 min to remove entrapped air.

### 2.3. Contact Angle Measurements

Contact angle (CA) measurement was performed to investigate hydrophobicity of custom silicones. To measure the CA first a custom goniometer setup was built similar to the existing literature [31]. After the samples were prepared, they were placed on the goniometer and 3 µl water was dropped on the surface and imaged. Images were analysed with Low-bond axisymmetric drop shape analysis (LBADSA) [32] method on ImageJ (National Health Instruments (NIH), Bethesda, MA, USA).

### 2.4. Mechanical Testing

Samples for durometer testing were prepared and measured in accordance with ASTM D2240-15(2021) [33]. Shore A hardness was determined using a Tekcoplus Shore A durometer (Hong Kong, China), and the results were reported as Hardness A (HA).

Compression test specimens were prepared and evaluated according to ASTM D395-18(2025) [34]. Formulations containing vPDMS/PMHS ratios of 7:3, 8:2, and 9:1 were cast into cylindrical molds. Following curing, the samples were tested to 50% strain or until failure using an Criterion model C43 electromechanical test system (MTS, Eden Praire, MN, USA) equipped with a 5 kN load cell. The measured values were converted into stress–strain data, and the initial elastic modulus was determined from the initial slope of the resulting curve within the 0–10% strain region.

### 2.5. Scanning Electron Microscopy Imaging

An 8:2 vPDMS:PMHS silicone was printed on nitrocellulose (NC) and cured, mounted on carbon tape, and sputter coated with iridium (15 nm; Leica EM ACE600). Surfaces were imaged on a Zeiss GeminiSEM 450 at 5 kV at magnifications of 300×, 400×, 2000×, and 5000× to visualize the NC, the NC–silicone interface, and the silicone surface.

### 2.6. Rheology Measurements

To perform rheological measurements a DHR-3 rheometer (TA instruments, New Castle, DE, USA) equipped with a 25mm parallel plate was used. All the measurements were obtained in room temperature (23 ºC). The samples were prepared with different silica filler concentrations and catalyst was excluded from the formulations to prevent crosslinking. The shear rate sweep was conducted between 0.1 and 100 s-1 shear rate. The thixotropic recovery sweep test was conducted by applying 0.1 and 100 s-1 shear rates with 60 s intervals. The amplitude sweep test was performed between 0.1 and 1000 Pa shear stress (or until rheometer reached its limits) at a frequency of 1 Hz. The frequency sweep test was performed between 0.1 and 100 rad s-1 angular frequency at 1% shear strain.

### 2.7. Fabrication of the Pneumatic Extrusion-based 3D Printer

A custom extrusion-based 3D printer was constructed based on an established design in the literature [35]. The summary of fabrication is as follows. First, a fused deposition modeling 3D printer (Adventurer 5M, Flashforge, Hangzhou City, Zhejiang, China) was obtained to be converted to the pneumatic 3D printer. The printhead was replaced with a custom one to house a 3 ml syringe barrel (Nordson, Westlake, OH, USA). Air pressure is controlled by solenoids and regulated with a high-flow precision compressed air regulator (McMaster-Carr, Elmhurst, IL, USA). Endstop switches and printbed adaptors were included in the 3D printer frame. To enable control, the microcontroller was replaced with creality motherboard V4.2.7 (Shenzen, Guangdong, China), rewiring was done and the firmware was updated with Marlin.

### 2.8. 3D printing of Silicone ink

The custom extrusion 3D printer was used to perform all the 3D printing experiments. To investigate the effects of printing parameters on filament diameter, silicone ink was printed with different needle sizes (22 G and 23 G), printing speeds (0.5, 1, 1.5, 2, 3, 4, 5, and 6 mm/s) and different pressures (100, 140, and 180 kPa). To measure printability of the inks, bilayer grids with increasing pore sizes were printed and printability index was obtained from analyzing the pore geometries [36,37].

### 2.9. Fabrication and Testing of LFA strips and µPADs

Each LFA strip consisted of a sample pad, nitrocellulose (NC) membrane, and absorbent pad assembled on an adhesive backing card with an overlap of ≈ 2 mm between adjacent components. Gold nanoparticle (AuNP)-conjugated with detection Mouse anti-pLDH Pan Malaria IgG antibody was prepared separately and stored at 4 °C until use. The NC membrane contained one test (T) line and one control (C) line. Capture Mouse anti-pLDH Pan Malaria IgG antibody and anti-mouse IgG antibody were dispensed onto the NC membrane to form the test and control lines, respectively, using an automated dispenser (Claremont Biosolutions, Upland, CA, USA). The membrane was dried at 37 °C for 1 hour. The distance between the test and control lines was maintained at 5 mm. The fully assembled strips measured ≈ 4 mm × 60 mm and were stored in sealed pouches at room temperature until further use.

To mitigate flow 2, three-sided, square-shaped silicone micro-barriers were printed on the NC membrane. The silicone barriers had a thickness of 0.5 mm and a width and length of 1 mm. The G-code for 3D printing was generated using MATLAB. The barriers were placed at distances of 2 and 3 mm from the edge of the sample pad. After printing, the membranes were placed in a 60 ºC oven for one hour. To measure the flow speed, strips were placed in a 96 well plate containing 120 µl of running buffer and distance travelled was measured from the video recording.

For both the conventional LFA and flow confined LFA, the *Pan Malaria pLDH* standards with concentrations of 100, 10, and 1 ng/mL were prepared in PBST buffer (PBS containing 1% BSA and 0.05% Tween 20). For the assay, the antigen samples were mixed with AuNP-conjugated with anti-pLDH Pan Malaria IgG antibody and incubated at room temperature for 15 minutes to facilitate immune complex formation. Subsequently, 120 µL of the reaction mixture was applied to the sample pad.

The sample migrated along the strip through capillary action. The antigen antibody AuNP complexes were captured at the test line by immobilized Mouse anti-pLDH Pan Malaria IgG antibody, generating a visible signal. Excess conjugates continued to migrate and were captured at the control line by anti-mouse IgG antibodies. After ≈ 15 minutes of incubation at room temperature, images of the strips were captured using the camera and the Quantitative analysis was performed using Image analysis software by measuring the test line intensity.

µPADs were fabricated in a similar fashion. Silicone barriers were printed on NC membranes using the custom 3D printer. The G-code for 3D printing was generated using MATLAB. After printing, the membranes were placed in a 60 ºC oven for one hour and colored water was used on µPADs demonstrate flow.

### 2.10. Statistical analysis

All data are presented as mean ± standard deviation. Statistical differences were determined by one-way analysis of variance (ANOVA) . Differences were considered statistically significant at p ≤ 0.05 (*), p ≤ 0.01 (**), and p ≤ 0.001 (***).

## 3. Results and Discussion

### 3.1. Development of extrudable hydrophobic silicone-based ink

We formulated a 3D-printable silicone-based ink comprising vinyl-terminated polydimethylsiloxane (vPDMS), polymethylhydrosiloxane (PMHS), dimethyldicholorosilane treated fumed silica filler (SF), a thixotropic agent, Karstedt’s catalyst, and 1-ethynyl-1-cyclohexanol inhibitor (**Fig. 1**). The vPDMS/PMHS pair enables platinum catalyzed hydrosilylation [38,39], filler and thixotropic agent tune shear-thinning behavior for extrusion. Catalyst/inhibitor ratios were adjusted to suppress premature curing for >24Lhr at room temperature yet permit crosslinking within 1□h at 60□°C.

**Fig. 1.**
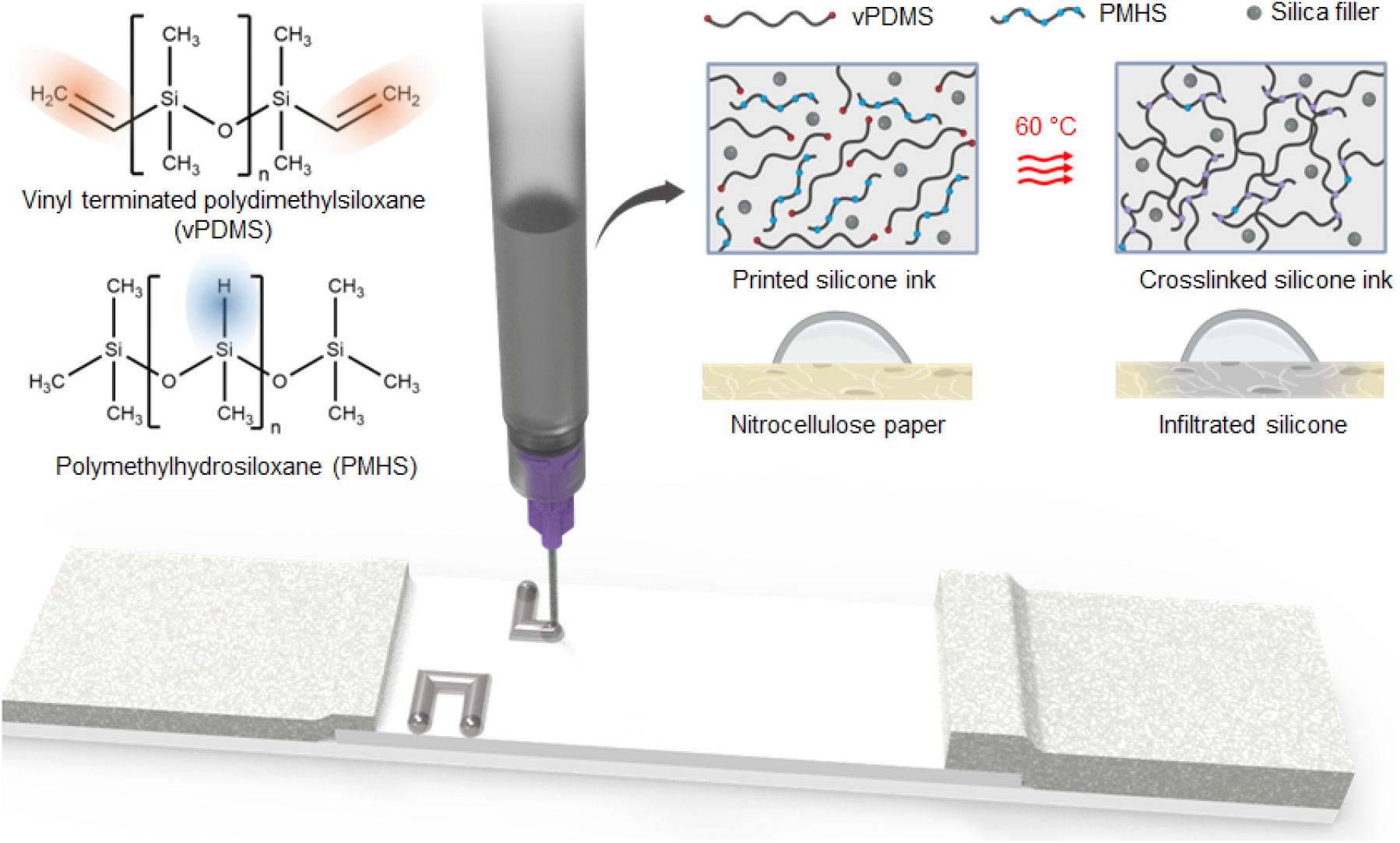
A schematic illustrating the contents and printing of silicone hydrophobic block.

### 3.2. Material characterization

Hydrophobicity of the cured silicone samples was evaluated using static water contact angle measurements. A droplet of deionized water was placed on the sample surfaces, and images were captured after droplet stabilization and analyzed with LBADSA technique (**Fig. 2a**) . The measured contact angles of all sample were around ∼110° (**Fig. 2b**) similar to reported values for other silicone [40], indicating the intrinsically hydrophobic nature of the silicone material. These results demonstrate that the developed silicone ink is suitable for hydrophobic blocks and other applications that would need hydrophobicity.

**Fig. 2.**
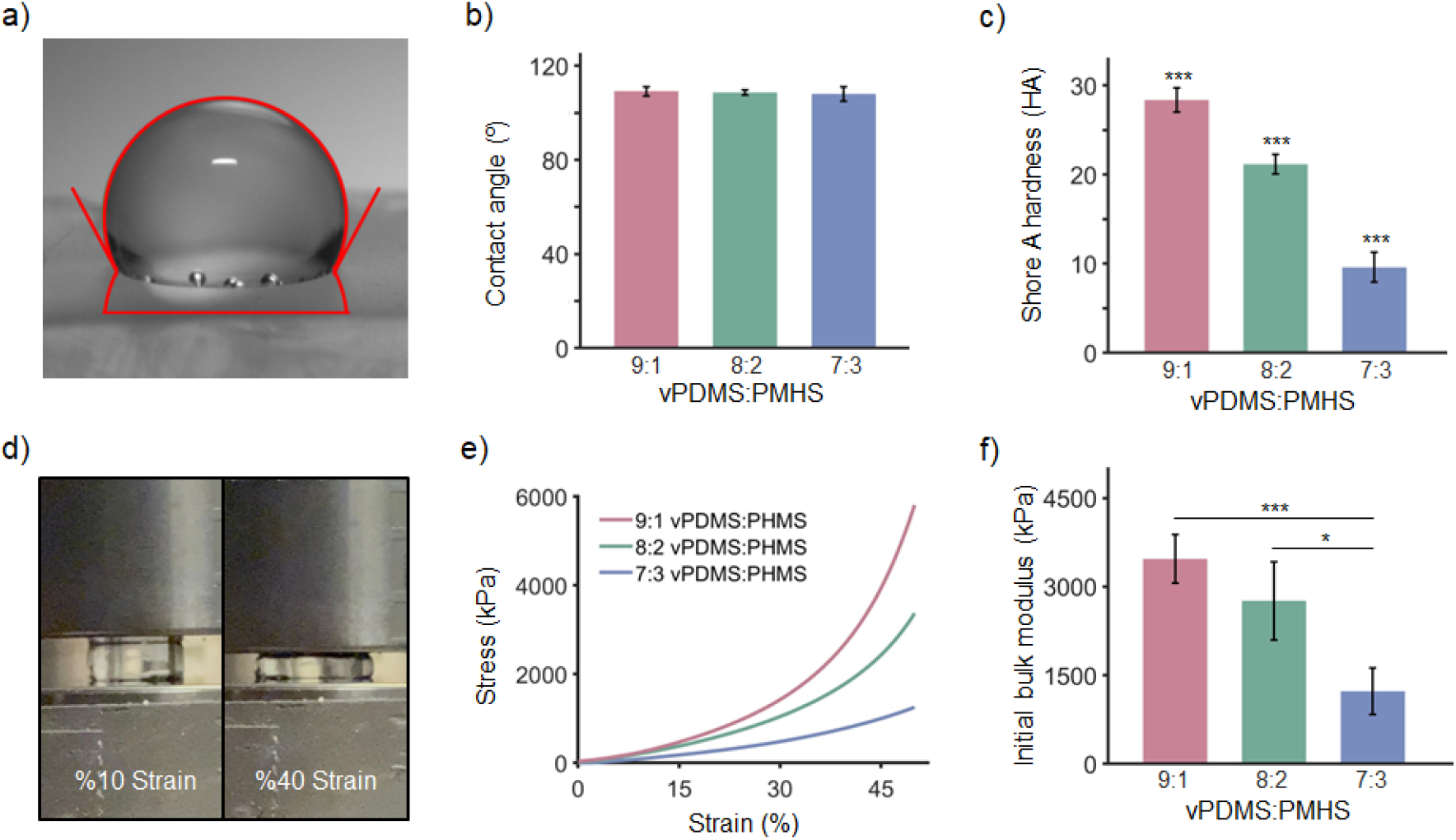
Material characterization of silicone ink. **a)** Contact angle calculation using a water droplet on silicone surface with LBADSA technique. **b)** Contact angles of different compositions of silicone formulations. **c)** Shore A Hardness of silicone compositions. **d)** Compression test. **e)** Compressive stress-strain curve. **f)** Initial bulk modulus of silicone compositions.

To understand mechanical strength of the silicone after its cured and to investigate how ratio between vPDMS and PHMS changes this strength, Shore A hardness test was applied to the samples (**Fig. 2c**). As the PHMS concentration increased and the vPDMS decreased, softer silicone was obtained. Overall, all tested formulations remained in the soft material regime. To evaluate the mechanical durability of the silicone ink, compression tests were performed on different silicone formulations (**Fig. 2d**). Most samples remained durable up to 50% compressive strain, and all formulations exhibited substantial resistance to applied stress (**Fig. 2e**). Consistent with the hardness test results, increasing the PMHS content produced softer silicone formulations (**Fig. 2f**).

To examine the surface topography of the printed samples, SEM imaging was performed (**Fig. 3 and Fig. S1**). Three distinct regions were imaged: the pristine NC paper, the interfacial region where silicone infiltrated the NC substrate, and the silicone surface. The images clearly demonstrate that the silicone penetrates and coats the NC paper.

**Fig. 3.**
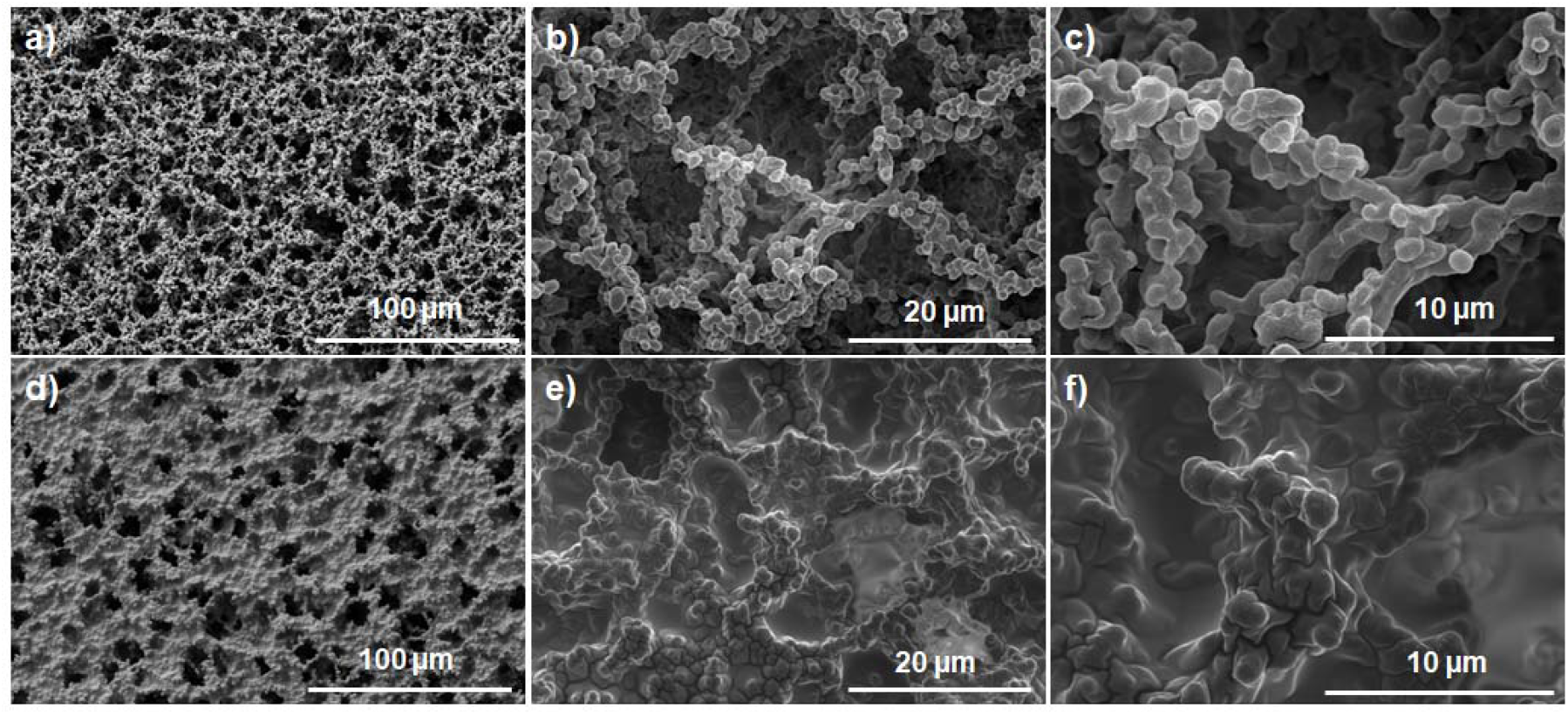
Surface topography visualization. SEM images of NC paper at **a)** 400x **b)** 2000x and **c)** 5000x magnification, silicone coated NC paper at **d)** 400x **e)** 2000x and **f)** 5000x magnification.

To achieve reliable flow confinement on nitrocellulose strips, we engineered and characterized a silicone-based hydrophobic ink optimized for 3D printing. The following rheological analyses ensured that the ink possessed appropriate flow, recovery, and printability characteristics. Rheological tests were conducted to assess the extrusion printability of silicone ink and to characterize its response under applied shear [41,42]. Also, the effect of SF concentration on the ink’s rheological properties was evaluated to analyze which concentrations can be used for an extrudable ink. First, a shear rate sweep was applied to inks with SF concentrations of 2, 4, 6, and 8% (w/w) (**Fig. 4a**). To describe the viscoelastic properties of the silicone material, the Herschel–Bulkley model was used [43,44], as given below:

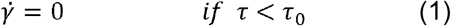

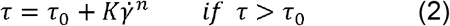

where 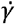 was the shear rate, *τ* was the shear stress, *τ*_0_ was the yield stress, K was the consistency index and n were the flow index. As SF concentration increased, stress and consistency of the ink increased significantly (**Fig. 4b**). The viscosity of all samples decreased with increasing shear rate (**Fig. 4c**). This behavior indicates shear thinning. Therefore, all formulations were suitable for extrusion applications. Also, as the concentration of SF increased viscosity increased as expected. Another important parameter for extrudable inks is self-recovery. After extrusion, the ink must rapidly regain its gel-like properties. To see the effect of SF concentration on self-recovery properties a 3-interval thixotropy test was applied to all concentrations (**Fig. 4d**). For all concentrations, inks displayed fast self-recovery and completely recovered within seconds. The viscoelastic behavior of silicone ink was evaluated using an amplitude sweep test (**Fig. 4e**). The result showed a linear viscoelastic region in which the storage modulus (G′) exceeded the loss modulus (G″) for all concentrations, indicating a stable gel-like state. At the flow point, where G′ and G″ intersected, the material transitioned from a gel to a liquid-like state. With further increases in shear stress, G″ became greater than G′, confirming viscous behavior. Results of the amplitude sweep are in line with the sheer rate sweep test and Hershey Buckley model. These findings demonstrate that the silicone ink yields and flows under applied shear stress, which is necessary for extrusion based 3D printing [41,45]. Frequency sweep tests were also conducted within the linear viscoelastic region to investigate the frequency dependence of silicone ink (**Fig. 4f**). Across the entire frequency range, all samples exhibited G′ > G″ expect %2 SF at high frequency, indicating stable gel like behavior.

**Fig. 4.**
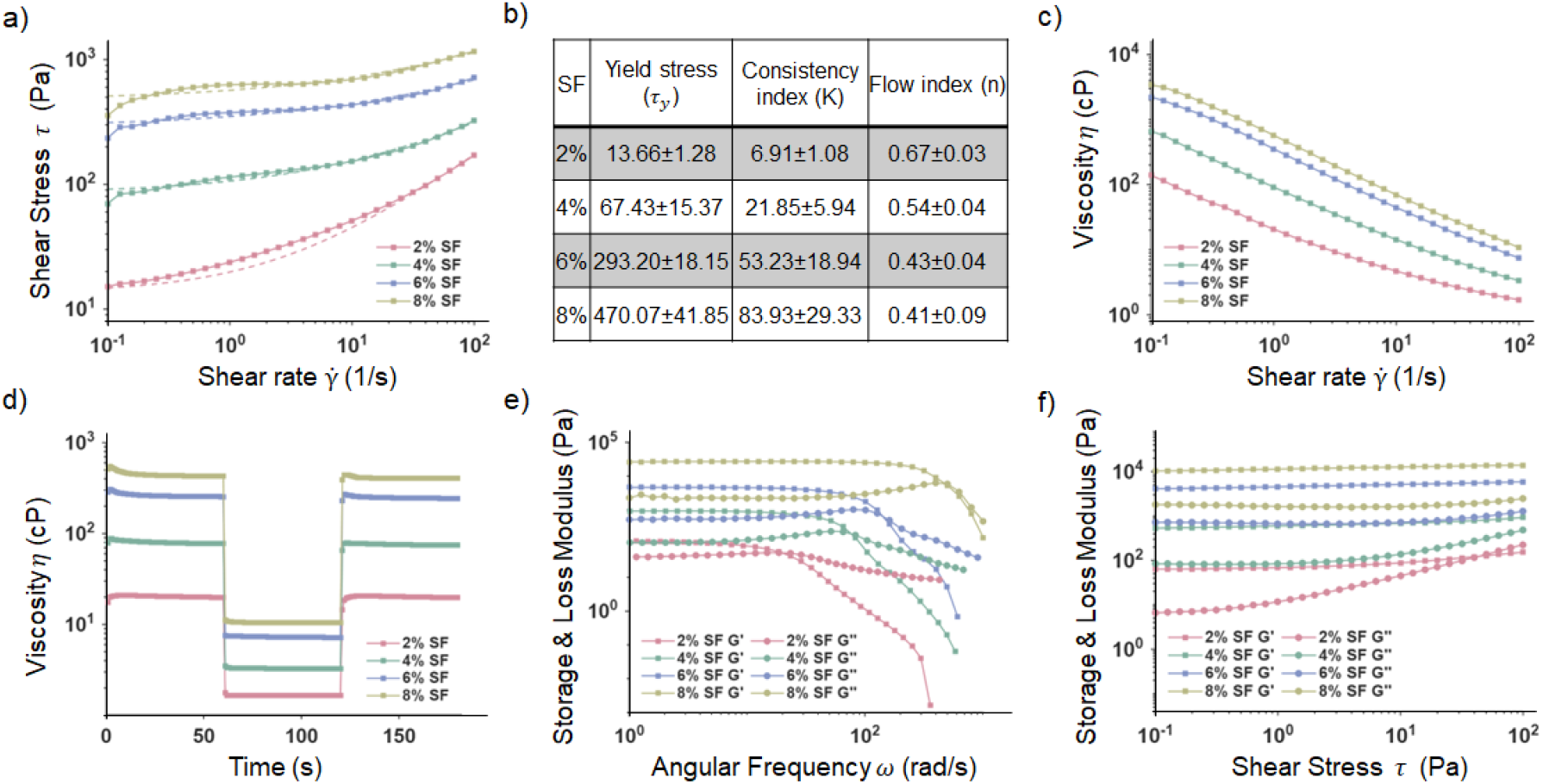
Rheological properties of silicone ink. **a)** Shear rate sweep test showing shear stress change under shear rate from 0.1 to 100 s-1, where dashed lines represent Hershel–Bulkley curve fittings. **b)** Hershel-Buckley model parameters obtained from shear rate sweep test. **c)** Shear rate sweep test showing viscosity change under shear rate from 0.1 to 100 s-1. **d)** 3-interval thixotropy test where the viscosity measured under alternating shear rates of 0.1 and 100 s-1. **e)** Amplitude sweep test showing response under shear stresses from 0.1 to 1000 Pa. **f)** Frequency sweep test showing response under angular frequencies from 0.1 to 100 rad s-1.

### 3.3. 3D printing of hydrophobic micro-barriers and print fidelity

To 3D print the silicone ink a custom, inexpensive pneumatic extrusion-based 3D printer was built (**Fig. S2**). The width of the printed blocks could be precisely controlled by adjusting the printing parameters (**Fig. 5a**) [46,47]. Increasing printing speed resulted in a reduction in filament width, while higher extrusion pressures led to an increase in filament width. By using a 22 Ga needle, we were able to obtain filament widths from ∼550 to 2700 µm by changing the pressure between 100 to 180 kPa (**Fig. 5b**). Within the same pressure range, we were able to obtain filament widths from ∼280 to 1700 µm with a 23 Ga needle (**Fig. 5c**). Filaments below this region resulted in printing failure. This level of tunability provides accurate control over feature dimensions, enables the fabrication of application specific complex printed structures [48,49]. We used printability index (Pr) as a quantitative metric to assess how accurately silicone ink was printed [36]. Where Pr < 1 represents filament spreading or over-extrusion, Pr=1 represents excellent print fidelity and, Pr>1 represents filament collapse, irregular shaped filaments or under extrusion. The Pr was calculated with the following formula:

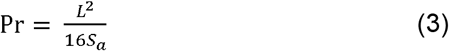

**Fig. 5.**
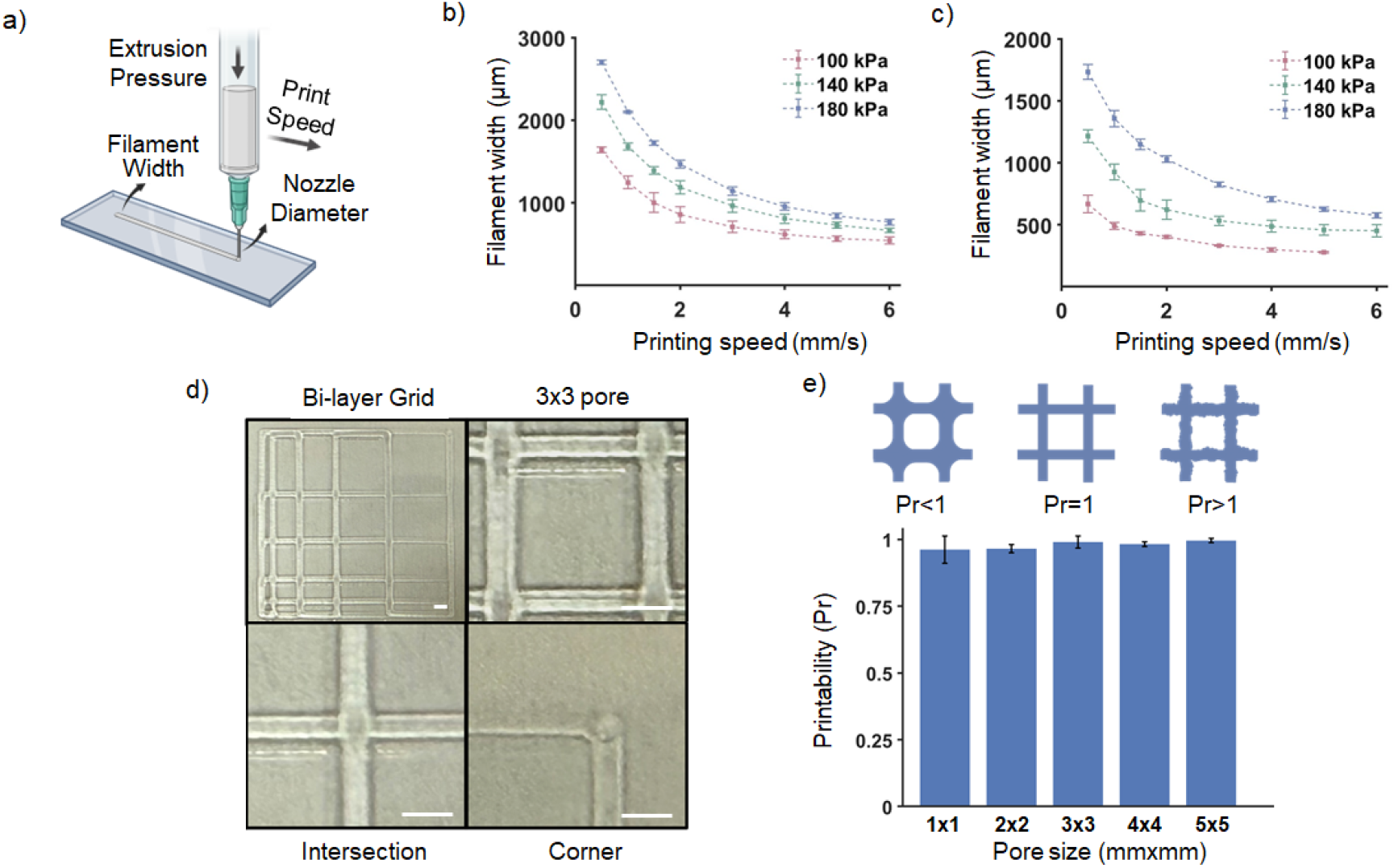
Printing characterization of silicone ink. **a)** A schematic displaying the process of extrusion printing and its parameters. Filament width at different print speeds and extrusion pressure for **b)** 22 G and **c)** 23 G needles. **d)** Bi-layer grid printed with silicone ink (scale bar represents 1mm). **e)** Printability measurements of the silicone ink

Where *L* was the perimeter and *S*_*a*_ was the area of the pore. The silicone ink exhibited satisfactory printability across all tested pore sizes (**Fig. 5d-e**). Collectively, these results demonstrate that the formulated silicone-based ink combines strong hydrophobicity, mechanical robustness, viscoelastic behavior, and precise extrusion printability, enabling the fabrication of well-defined micro-barrier structures. These features ensure reliable flow confinement within lateral flow strips, providing the foundation for the enhanced analytical performance of the silicone block assisted LFA system.

### 3.4. Development of LFAs and µPADs with hydrophobic micro-barriers

Uncontrolled lateral fluid dispersion within the NC membrane can limit test line signal intensity, particularly at low analyte concentrations [50,51]. To address this, we utilized the hydrophobic extrudable silicone-based ink to fabricate micro-barrier structures that guide fluid flow and enhance analytical sensitivity (**Fig. S3**). The printed hydrophobic silicone micro-barriers confine lateral flow and moderately slow analyte migration, thereby increasing residence time at the test zone. Time-resolved wicking images and quantitative flow analysis (**Fig. 6a-c**) show that barrier integration regulates capillary transport, reducing lateral dispersion compared to pristine NC and shown as a (**Video 1**). The performance of the flow-confined NC membrane incorporating 3D-printed silicone micro-barriers was evaluated against a pristine NC membrane representing a conventional LFA format using Pan-malaria pLDH antigen standards at concentrations of 100, 10, and 1 ng/mL. In the conventional LFA, the test line was strongly visible at 100 ng/mL, weakly detectable at 10 ng/mL, and nearly absent at 1 ng/mL. In contrast, the flow-confined LFA generated substantially stronger and more uniform test-line signals across all tested concentrations (**Fig. 6d**). Importantly, the 1 ng/mL antigen concentration, which approached the detection threshold of the conventional strip, produced a clearly visible and quantifiable signal in the flow-confined configuration. Quantitative image analysis further confirmed the enhanced analytical performance of the flow-confined design. This enhancement effectively lowered the practical detection limit from 10 ng/mL in the conventional format to 1 ng/mL in the flow-confined system, corresponding to a ten-fold improvement in analytical sensitivity (**Fig. 6e**).

**Fig. 6.**
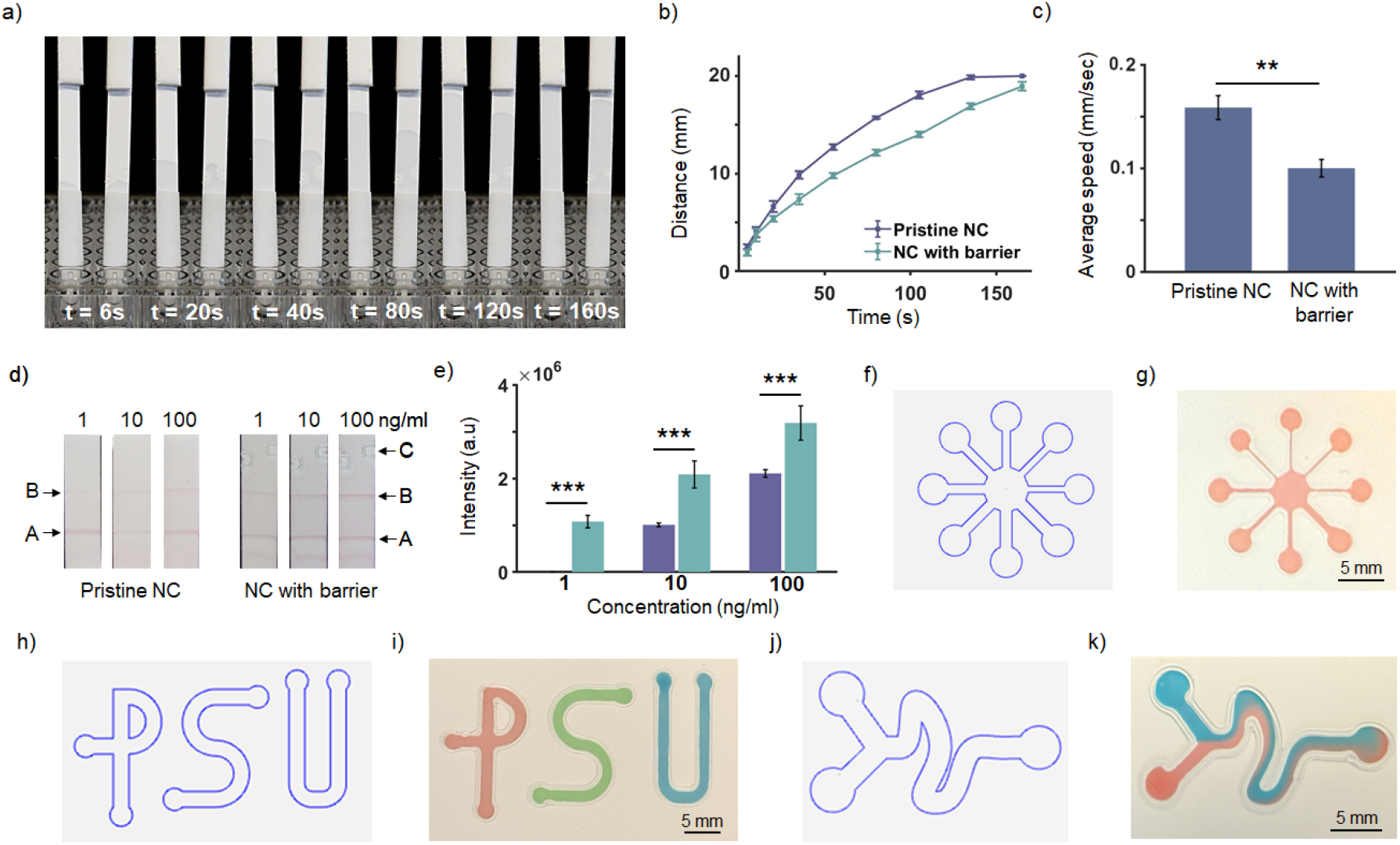
Fabrication of LFAs and µPADs with hydrophobic micro-barriers. **a)** Wicking speed test for pristine NC membranes and NC membranes with silicone micro-barrier **b)** Wicking distance over time **c)** Average flow speed. **d)** Representative strips with increasing concentrations of pLDH antigen (A: control line, B: test line, C: silicone micro-barriers). **e)** Measured test line intensity. Radiating and varying micro-channels **f)** design and **g)** print. “PSU” micro-channels **h)** design and **i)** print. Flow focusing mixer **j)** design and **k)** print.

Overall, these findings demonstrate that simple geometric modification of NC membranes using 3D-printed silicone micro-barriers can significantly improve LFA performance without requiring changes to the assay biochemistry. The ability to detect pLDH at concentrations as low as 1 ng/mL with strong visual clarity highlights the potential of flow-confined LFAs for improved malaria diagnosis, particularly in low-parasitaemia or early-stage infections where conventional rapid diagnostic tests often exhibit limited sensitivity. Because the barrier printing strategy is compatible with existing LFA manufacturing processes and utilizes low-cost silicone inks with scalable extrusion-printing methods, this approach offers strong potential for translation into practical point-of-care diagnostic platforms.

To further demonstrate the capabilities of this technique, we developed several distinct µPAD designs with varying geometries and fluidic layouts. Radiating and varying microchannel (**Fig. 6f-g, Video 2**), “PSU” microchannels (**Fig. 6h-i**), and a flow-focusing mixer (**Fig. 6j-k**) were also designed and fabricated to demonstrate the versatility of the fabrication approach. The successful formation of these patterns highlights the adaptability of the method for producing customized paper microfluidic platforms with precise fluid pathways. This design flexibility is particularly important for expanding the functionality of µPADs toward more complex analytical applications, including controlled sample distribution, multiplexed detection, and integrated assay formats. We also evaluated the minimum channel dimensions capable of supporting fluid flow in the fabricated devices. Using hydrophobic barriers with a width of 300 µm, we successfully achieved fluid flow through a microchannel with a width of 150 µm (**Fig. S4)**. These results further demonstrate the potential for the fabrication of compact, intricate and high-resolution paper microfluidic devices.

## 4. Conclusion

In conclusion, we report a silicone extrudable ink platform for the fabrication of high-resolution hydrophobic features on paper substrates. By formulating a vPDMS/PMHS system and tailoring its rheological behavior with hydrophobically treated silica fillers and a thixotropic agent, we obtained a shear-thinning ink capable of controlled extrusion and stable feature formation. This strategy enables simple, low-cost, rapid, and robust patterning of hydrophobic barriers while overcoming the limited resolution typically associated with unmodified liquid PDMS. Due to the chemical and thermal inertness, mechanical durability, and long-term stability of silicone, the resulting barriers are well suited for reliable paper-based diagnostic applications. Importantly, the developed platform enabled the fabrication of high-resolution paper microfluidic devices down to 150 µm channel width and improved the sensitivity of a lateral flow assay for malaria detection. Overall, this work establishes a practical and scalable strategy for advancing paper-based microfluidics and diagnostic technologies.

## Acknowledgments

The authors acknowledge the support of Pennsylvania State University in conducting this work. D.P. acknowledges funding from the Centers for Disease Control and Prevention (75D30122C15492), the National Science Foundation (CBET 2153091 and 2229986). Authors would like to thank Yunzhen Zheng for helping with the SEM imaging. Dr. Federico Harte and Dr. Taha Ahmed for proving access and help for the rheometer. Multiple figures were generated using BioRender.com.

