## Supplementary Information for "Extrusion-Printed Silicone Microarchitectures for Geometry-Controlled Flow in Lateral Flow Diagnostics and Paper Microfluidics"

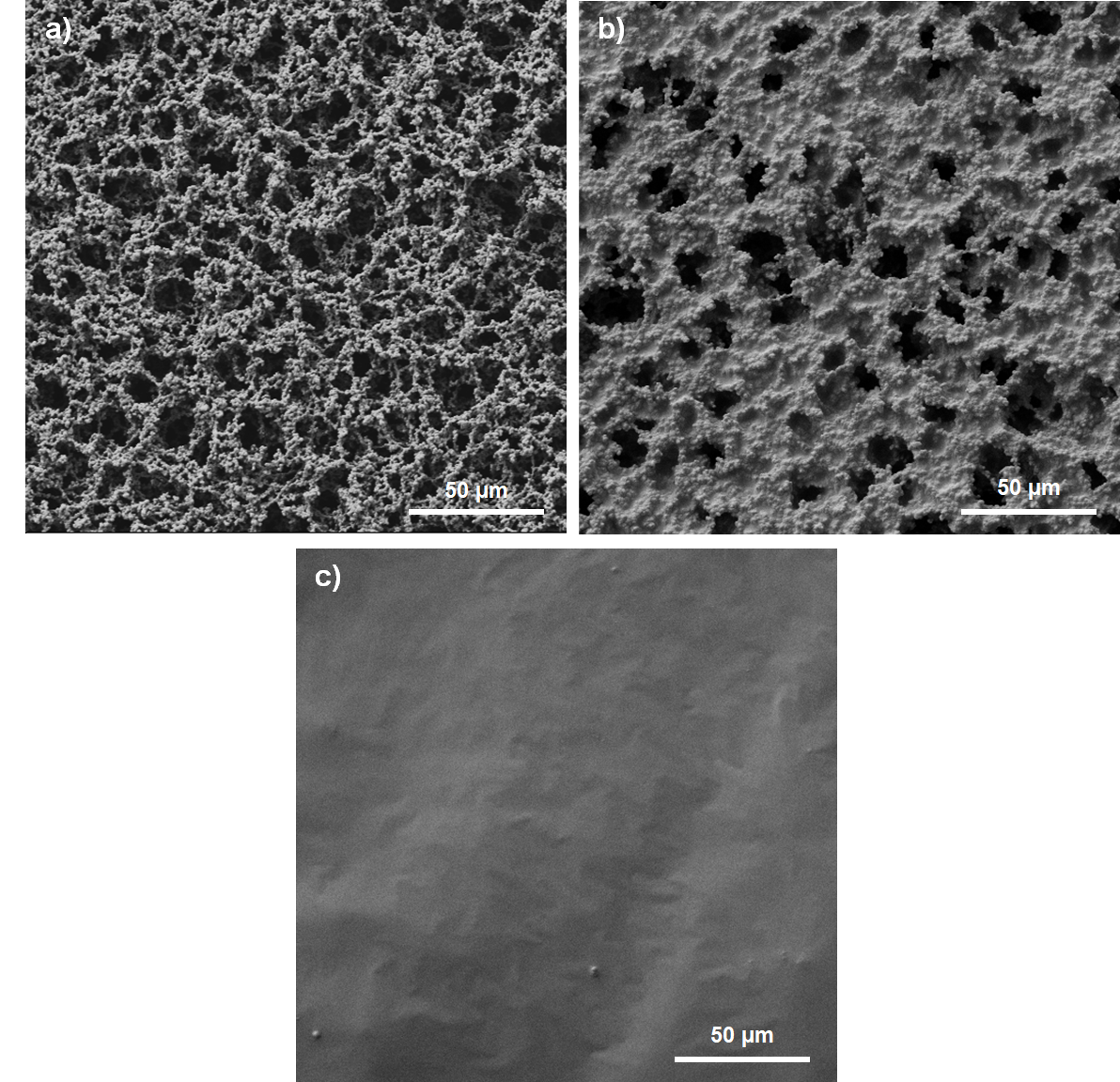

**Fig. S1**. SEM figures of **a)** pristine NC paper, **b)** silicone coated NC paper and **c)** printed silicone at 300x magnification.

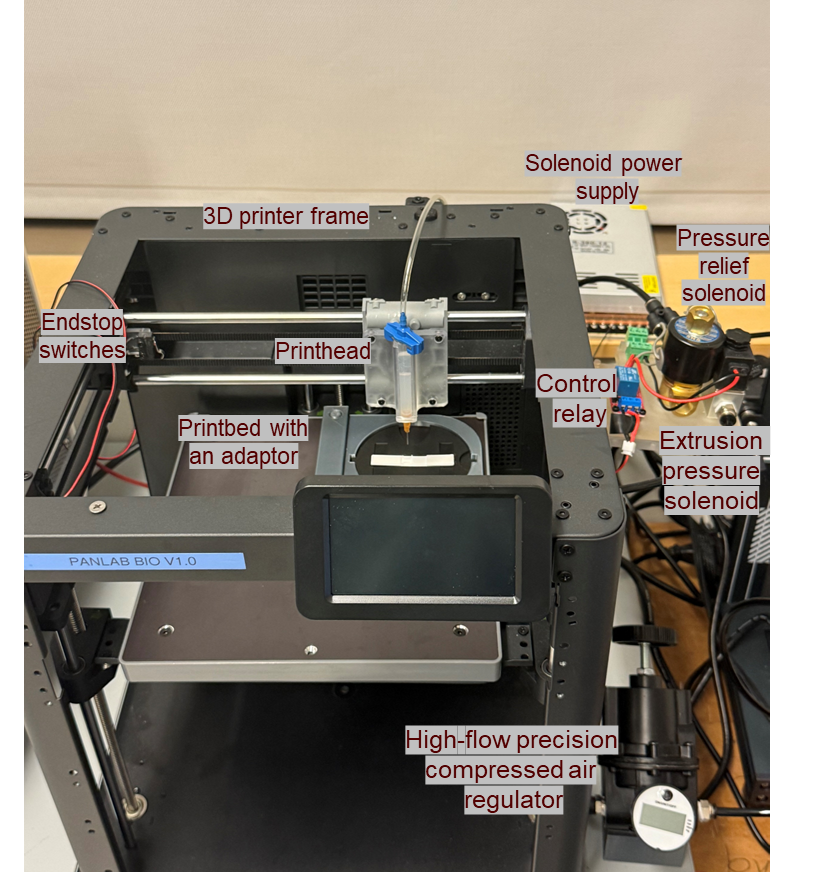

**Fig. S2**. Custom built pneumatic extrusion-based 3D printer

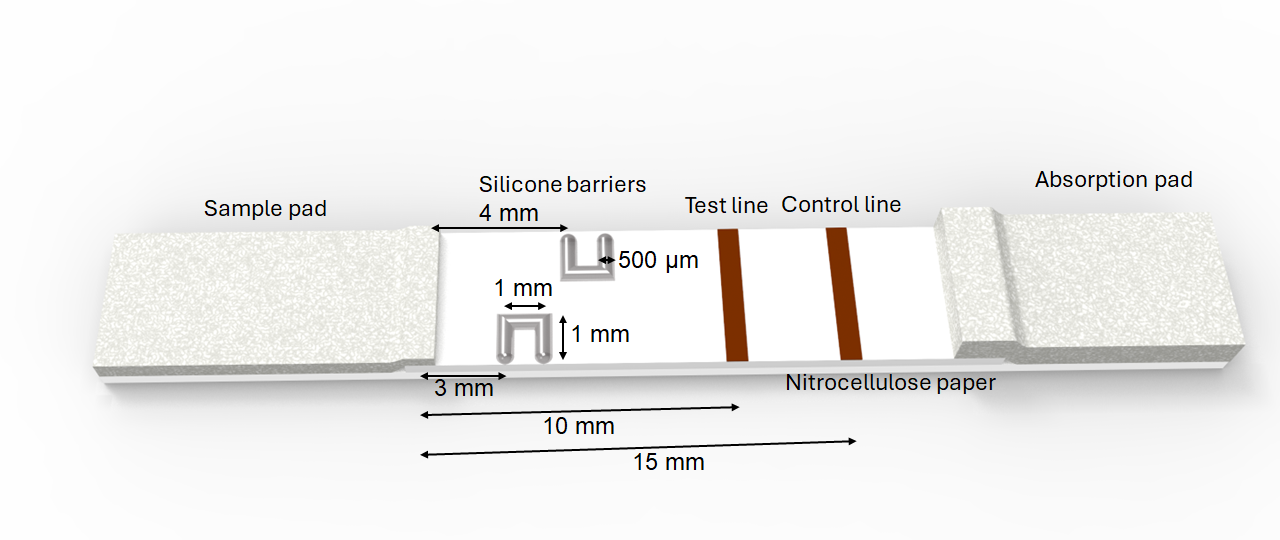

**Fig. S3.** Schematic illustration of the lateral flow strip integrated with silicone micro-barriers showing channel geometry and dimensions (total length 15 mm; reaction zone 10 mm; barrier spacing and widths as indicated, including 500 µm features)**.**

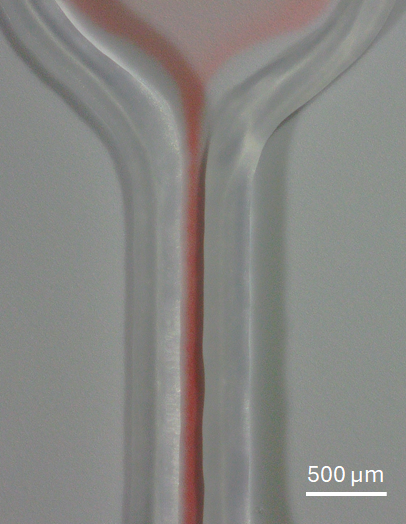

**Fig. S4.** Microfluidic channel with 150 µm width
